# Ton Motor Conformational Switch and Peptidoglycan Role in Bacterial Nutrient Uptake

**DOI:** 10.1101/2023.08.11.552980

**Authors:** Maximilian Zinke, Maylis Lejeune, Ariel Mechaly, Benjamin Bardiaux, Ivo Gomperts Boneca, Philippe Delepelaire, Nadia Izadi-Pruneyre

## Abstract

Active nutrient uptake is fundamental for survival and pathogenicity of Gram-negative bacteria, which operate a multi-protein Ton system to transport essential nutrients like metals and vitamins. This system harnesses the proton motive force at the inner membrane to energize the import through the outer membrane, but the mechanism of energy transfer remains enigmatic. Here, we study the periplasmic domain of ExbD, a crucial component of the proton channel of the Ton system. We show that this domain is a dynamic dimer switching between two conformations representing the proton channel’s open and closed states. By *in vivo* phenotypic assays we demonstrate that this conformational switch is essential for the nutrient uptake by bacteria. The open state of ExbD triggers a disorder to order transition of TonB, enabling TonB to supply energy to the nutrient transporter. We also reveal the anchoring role of the peptidoglycan layer in this mechanism. Herein, we propose a mechanistic model for the Ton system, emphasizing ExbD duality and the pivotal catalytic role of peptidoglycan. Sequence analysis suggests that this mechanism is conserved in other systems energizing gliding motility and membrane integrity. Our study fills important gaps in understanding bacterial motor mechanism and proposes novel antibacterial strategies.

## INTRODUCTION

Gram-negative bacteria present a unique challenge for the development of novel drugs due to their dual-membrane structure, which effectively protects them by preventing many antibiotics accessing their targets within the cell.^1^ This dual-membrane architecture requires specialized transport systems for essential nutrients – like iron, nickel, vitamin B12 and certain carbohydrates^2^. These gram-negative specific systems guarantee efficient transport over both the inner and the outer membrane as well as the periplasm, potentially creating vulnerabilities for therapeutic intervention. One of the key systems involved in this transport is the Ton system, which forms a multi-protein complex embedded in the inner membrane (Figure 1a). This system utilizes the proton motive force (PMF) to physically open a variety of outer membrane transporters – the so called TonB-dependent transporters (TBDTs) – and, hereby, realizes active transport by means of the inner membrane proton channel forming complex ExbB-ExbD and the periplasm crossing protein TonB. While the TBDTs are nutrient-specific, TonB and ExbB-ExbD can be employed to multiple TBDTs (multi-target).^3^ In addition to this multi-target Ton system, some Ton systems are dedicated to a specific transporter (single-target). This is the case of the Heme Acquisition System (Has) allowing bacteria to acquire heme as an iron source and involving orthologs of ExbB, ExbD with a TonB paralog called HasB.^4–6^

**Figure 1.**
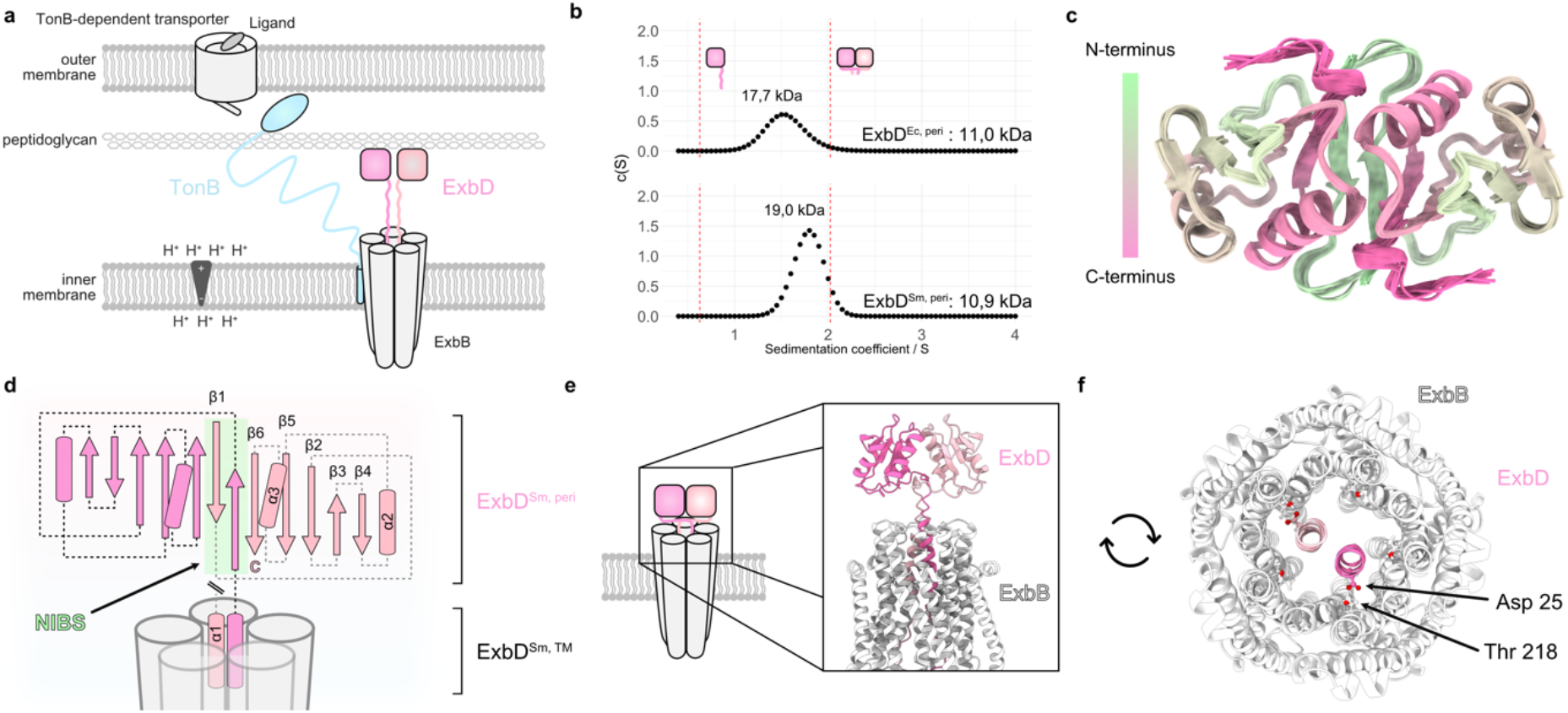
The periplasmic domain of ExbD forms a homodimer. **a** The Ton system consists of an inner membrane proton channel – formed by ExbB (grey) and ExbD (pink) – and the periplasm spanning protein TonB. (blue). The latter links the system to a TonB-dependent transporter in the outer membrane. The globular C-terminal domain of ExbD (pink boxes) is connected to its helical N-terminus, which is inserted into the ExbB channel, by a disordered region (pink, wiggly lines). **b** Sedimentation coefficient distribution obtained from analytical ultracentrifugation of ExbD^Sm, peri^ and ExbD^Ec, peri^ indicates that both proteins exist in a dimeric state at physiological pH. The red dashed lines represent estimated sedimentation coefficients for monomeric ExbD and dimeric ExbD as depicted by the cartoons (pink). The theoretical molecular weights of both species are indicated above. **c** The top view of the NMR structure ensemble of dimeric ExbD^Sm, peri^ reveals that the N-terminal residues 44-49 (green) form an intermolecular β-strand. **d** Homodimeric ExbD^Sm, peri^ consists of two monomers (pink and light pink), each with a β-sheet (arrows) and two α-helices (barrels). Part of the dimeric interface is formed by a swapped, intermolecular and anti-parallel β-sheet called the N-terminal Intermolecular Beta-Strand (NIBS, green). The NIBS, composed of residues 44-49, connects the intramolecular β-sheets, creating a continuous β-sheet across the dimer interface. In the full-length protein, ExbD^Sm, peri^ is N-terminally connected to two α-helices (α1) that are embedded in the ExbB channel (grey). **e** Side view and **f** Top view of the AlphaFold2 model of the full ExbB-ExbD complex, generated using the NMR structure (PDB ID 8PEK, this work) as a template. In this model, the dimeric organization of the periplasmic domain of ExbD (pink and light pink) imposes the alignment of its helices in similar positions and orientations, and locking both Asp 25 sidechains in hydrogen bonds with Thr 218 from ExbB. In the top view, the periplasmic domain of ExbD is not visible due to its location above the clipping plane.

At the inner membrane, the Ton system consists of the proton-channel forming complex ExbB-ExbD and the protein TonB, which are together capable of harnessing the PMF. The structure of the whole complex is unknown. Recent cryo-EM data showed two α-helices situated in a periplasm-facing central pore of the pentameric ExbB channel. These densities were attributed to the most N-terminal part of an ExbD dimer (residues 12-42).^7^ However, no information about the organization of the C-terminal, periplasmic part of ExbD could be gained as it remains invisible in all known cryo-EM structures to date most likely due to its dynamics.^5,8,9^ The only structural data is a monomeric, solution NMR structure of the periplasmic domain of ExbD at pH 3 that features an unfolded N-terminus (residues 44-63) followed by a globular fold (residues 64-132).^10^ It has been proposed that a PMF-dependent de- and reprotonation of the highly conserved ExbD residue Asp 25 in the helix that is embedded in the pore of the ExbB channel, causes a rotation of ExbD within ExbB^11^ – analogous to the MotA-MotB system involved in flagellar motility.^12^ At the outer membrane, ligand binding to a TBDT results in the propulsion of an N-terminal extension of the TBDT into the periplasm, where it is tightly bound to TonB via a conserved region called the TonB box.^13^ TonB consists of an N-terminal α-helix anchoring it into the inner membrane and proposedly to ExbB^5^, a periplasm-spanning intrinsically disordered linker, and a C-terminal, globular domain responsible for TBDT binding.^14^ The intrinsically disordered linker contains a proline-rich region, approximately 70 residues in length, with an N-terminal stretch of Glu-Pro repetitions and a C-terminal stretch of Lys-Pro repetitions.^15^ These stretches are organized in polyproline II helices and behave like two static rods, which however still retain some flexibility.^16,17^ It is proposed that following TonB’s binding to the TonB box, the hypothesized ExbD rotation would generate a pulling force on TonB which unfolds a plug in the TBDT, opening a channel and ensuring selective ligand entry.^18^ These dynamics might vary in the Has system due to a couple of distinguishing features: 1.) The disordered linker of HasB does not contain an extensive stretch of Glu-Pro repetitions, consequently lacking two large regions of contrasting charges. 2.) HasB is likely perpetually bound to the TBDT, even in the absence of the ligand. This characteristic can be attributed to the single-target nature of the HasB, eliminating the necessity to accommodate multiple TBDTs.^19^

Due to the partial localization of the Ton system in the periplasm, a potential role of the periplasmic peptidoglycan layer within the mechanism of the system has been proposed.^20^ In the related MotA-MotB system, MotB contains a C-terminal peptidoglycan-binding motif that anchors the MotAB stator to the cell wall^21^, while in the analogous Tol-Pal system^22^, this interaction is facilitated by Pal and TolR^23,24^. However, the nature of this interaction in the Ton system has remained elusive.

Despite being first identified in 1978^25^, key steps of the Ton system’s mechanism remain poorly understood primarily due to incomplete structural information especially on the connection between ExbD and TonB. This is likely due to the disordered and dynamic nature of the periplasmic domains of both proteins, which renders them unsuitable for cryo-EM studies. Uncovering the role and the assembly of these two proteins is of great importance, as it could not only improve our understanding of nutrient transport in gram-negative bacteria but also provide new targets for antibiotics development. To ensure the broad relevance of our results, we investigated two representative systems: the multi-target Ton system from *Escherichia coli* and the single-target Has system from *Serratia marcescens*. These systems were chosen due to the plethora of existing functional data and a partial knowledge of their structures, allowing their comprehensive analysis. In this study, we use NMR spectroscopy to make the dynamic visible: we present the dimeric structure of the periplasmic domain of ExbD in different states, including a sparsely populated one. We demonstrate that this minor state is conformationally selected upon binding to an intrinsically disordered region (IDR) of TonB, which undergoes a disorder-to-order transition. Moreover, mutagenesis and *in vivo* phenotypic assays confirm that this multi-state transition of ExbD is required for its function. Also, we show that the transient interaction of the ExbD dimer to the peptidoglycan layer in the periplasm is crucial for the action of the Ton system as it is transiently bound by the ExbD dimer.

## RESULTS

### The periplasmic domain of ExbD forms a homodimer

We recombinantly produced the periplasmic domains of ExbD from both organisms (from *E. coli* including residues 43-141: ExbD^Ec, peri^, from *S. marcescens* including residues 43-140: ExbD^Sm, peri^; all constructs and NMR samples are summarized in Table S1-2). Analytical ultracentrifugation (AUC) revealed that the periplasmic domain of ExbD of both systems is organized as a dimer in solution (Figure 1b).

We proceeded our structural study with ExbD^Sm, peri^ as NMR fingerprint spectra indicated that ExbD^Ec, peri^ suffers from line broadening due to conformational exchange dynamics, while the *S. marcescens* protein revealed high-quality spectral data (Figure S1-S2). The solution structure of ExbD^Sm, peri^ was determined as detailed in the Methods and resulted in a swapped homodimer (Figure 1c and Table S3). Each protomer is composed of a four-stranded β-sheet on one side and two α-helices on the other (Figure 1d). The dimeric interface is established through i) a network of hydrophobic interactions centered around the sidechain of Tyr 112, and ii) a swapped, intermolecular and anti-parallel β-sheet, referred to as the N-terminal Intermolecular Beta-Strand (NIBS). The NIBS, formed by residues 44-49, links the β-sheets of each protomer creating a continuous, intermolecular β-sheet. This structure exhibits structural similarity with a crystal structure obtained for the periplasmic domain of TolR, the ExbD counterpart in the Tol-Pal system.^24^ However, this dimeric organization has not been observed in the only known ExbD^Ec, peri^ structure that was shown to be monomeric.^10^ This structure was solved at pH 3 and its NIBS region is disordered. Here, our experiments were performed at pH 7, and the ordered nature of the NIBS was also confirmed by relaxation measurements, as shown in Figure S3.

To contextualize the dimeric structure of ExbD^Sm,peri^ within the whole assembly, we used our solution structure as a template to generate a model of the full ExbB-ExbD complex with AlphaFold2 (Figure 1e, f).^26^ In the published cryo-EM structures, the ExbB-embedded helices of ExbD are not equivalent, displaying differences regarding their height and rotation towards each other as they are shifted by half a helical turn. Also it has to be noted, that in these structures only one of the two Asp 25 of ExbD is involved in hydrogen bond formation, leaving the other Asp 25 proposedly “ready” to accept a new proton for conduction.^5,7^ However, in the generated model with a dimeric organization of the periplasmic domain of ExbD, the helices are aligned in similar positions and orientations, directing both Asp 25 sidechains towards Thr 218 with a distance compatible with the formation of a hydrogen bond. Based on these observations, we postulate that this model, featuring both Asp 25 sidechains engaged in hydrogen bonds, could represent an inactive, proton-impermeable state of the channel.

### ExbD samples different conformational states

The absence of the periplasmic domain of ExbD in the cryo-EM reconstructions indicates that the dimer undergoes extensive conformational dynamics and does not simply occupy a single state as the NMR structure might imply. To characterize these conformational dynamics and to understand how the protein can fulfill its function, we explored the dynamical landscape of the periplasmic domain of ExbD. For this purpose, we analyzed ExbD^Sm, peri^ by ^15^N chemical exchange saturation transfer (CEST) experiments, which allow for the characterization and quantification of different exchanging conformational states of a protein – with populations as sparse as less than 1 % – that are in slow exchange (lifetimes ranging from ∼5-50 ms).^27^ The per-residue CEST profiles for the NIBS residues are characteristic of an exchange between three conformational states with a major state A (large dip) and two minor conformational states B and C (smaller dips) (Figure 2a and Figure S4).

**Figure 2.**
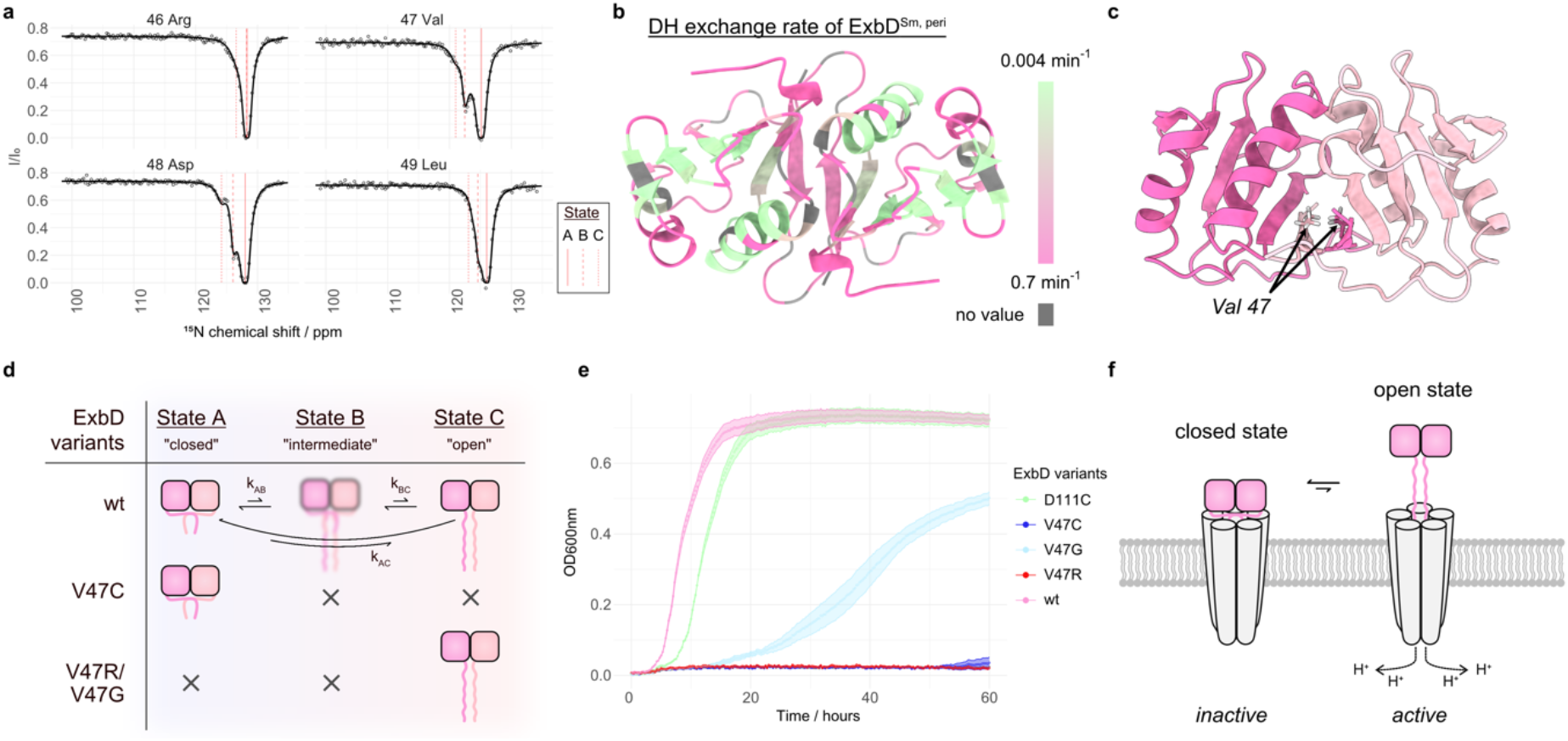
The periplasmic domain of ExbD samples different conformational states. **a** ^15^N-Chemical exchange saturation transfer (CEST) profiles reveal a 3-state exchange of the NIBS in ^15^N-labeled ExbD^Sm, peri^. The experimental data points and the least-square fit are represented by dots and solid lines, respectively. The fitted chemical shift values for states A, B, and C are indicated by red lines. Intensity ratios of NMR signals from residues 46 to 49 are shown. **b** Deuterium hydrogen (DH) exchange rates measured by NMR spectroscopy mapped onto the structure of ExbD^Sm, peri^ (fast exchange: pink, slow exchange: green, no value: grey) show that the NIBS undergoes faster amide proton exchange than adjacent β-strands, supporting the disordered nature of state C. **c** Side view of dimeric ExbD^Sm, peri^ where the monomeric units are colored in pink and light pink. The residue Val 47 is located at the dimer interface in the closed state. **d** Wild type (wt) ExbD^Sm, peri^ (pink) samples a major state A, and the minor states B and C. State A is characterized by a folded NIBS, while state C features a disordered NIBS. State B represents an intermediate state between these two extremes. The V47C mutant forms a disulfide bridge that locks ExbD^Sm, peri^ in the closed state, while the V47R and V47G mutants mainly yield the open state. **e** Bacterial growth curves of ExbD^Sm, peri^ variants wt (pink), V47R (red), V47G (light blue), V47C (blue) and the control D111C (green). Under *in vivo* conditions that necessitate the activity of the Has system, these mutations substantially impact bacterial growth, emphasizing that not merely the individual states of ExbD, but the exchange between them, is vital for the system’s proper functioning. **f** In a schematic representation of the ExbB-ExbD complex, the ExbD dimer exchanges between a closed, main state and a sparsely populated open state. The transition between the states is characterized by the un- and refolding of the NIBS residues. Herein, only the open state might be permeable to protons.

We utilized the software package ChemEx^28^ to fit the profiles and extract exchange parameters, including the chemical shifts of the minor conformations (Table S4). Intriguingly, upon re-examination of 2D TROSY spectra, we were able to identify the resonances for state C as low signal-to-noise peaks (Figure S5). Additionally, deuterium-hydrogen exchange spectroscopy of ExbD^Sm, peri^ revealed that the NIBS is more accessible to the solvent than the adjacent β-strands (Figure 2b and Figure S6). This observation, in conjunction with the chemical shifts of state C, indicates that state C is an “open” state characterized by the disordered nature of the NIBS residues, as opposed to the “closed” state A, where those residues form an intramolecular β-sheet, as shown in the NMR structure. It is important to note that the dimeric structure of ExbD^Sm, peri^ is maintained during this opening process. We hypothesize that state B marks an intermediate state between those extrema, as its chemical shifts lie in all cases in between those of state A and state C. We note that a similar conformational exchange process is present in ExbD^Ec, peri^; however, for this species, the exchange resides in the intermediate NMR timescale (μs-ms), as observed in the fingerprint spectra (Figure S2).

To confirm the exchange driven order-to-disorder transition of the NIBS, we designed Val 47 mutants for both ExbD^Sm, peri^ and ExbD^Ec, peri^. In the structure of ExbD^Sm, peri^ wild type (wt), this residue at the dimer interface occupies a hydrophobic pocket and is in close spatial proximity to the corresponding Val 47 from the other protomer within the homodimer (Figure 2c). The mutant V47C is thought to form a disulfide bridge locking the subunits together and, hence, mimicking a pure closed state. Conversely, V47R and V47G both would lead to a dimer with an unfolded NIBS as the arginine sidechain cannot occupy the hydrophobic pocket and the glycine perturbs the stability of the NIBS – resulting in a mainly open state (Figure 2d). These behaviors were confirmed by NMR spectroscopy, which can be briefly summarized as follows: a) The V47C mutation of ExbD^Sm, peri^ shows only small chemical shift perturbations in comparison to the wt, indicating that the main conformation of ExbD^Sm, peri^ is indeed the closed state. b) The V47C mutation of ExbD^Ec, peri^ results in NMR spectra absent of conformational exchange. c) The V47R and V47G mutants of both ExbD^Sm, peri^ and ExbD^Ec, peri^ exhibit NIBS residues with chemical shifts indicative of a disordered state (Figure S7-9).

Having established ExbD mutants that mainly occupy either the open (V47R/V47G) or closed state (V47C), we proceeded to study the impact of these mutations on the activity of the Has system *in vivo*. For that purpose, we monitored the growth of the Ton system-deficient *E. coli* strain *K12* C600*ΔhemAΔtonBΔexbBD*, which requires external heme as an iron source for growth. This deficiency was compensated by two plasmids, one encoding for the entire Has operon (hasISRADEB) from *S. marcescens* and a second one for ExbB-ExbD^Sm^. This setup allowed for a direct correlation of bacterial growth and the efficiency of heme import and, hence, reflecting the correct functioning of the Has system. We also verified the assembly of the wt and mutant ExbB-ExbD complexes and confirmed that the proteins are expressed at comparable levels in the membrane, ensuring that the observed effects on bacterial growth are attributable to the specific mutations and their impact on the ExbD conformational exchange (Figure S10). Consequently, we could analyse the influence of ExbD adopting exclusively the open or closed state on the functioning of the Has system. Compared to the wt and the control mutant D111C (a residue far from the NIBS), V47C and V47R completely abolish growth, while V47G significantly reduces growth (Figure 2e). These results demonstrate that neither the open state nor the closed state of ExbD alone, but rather the exchange between them, is crucial for the mechanism of the Has system, potentially reflecting distinct steps in the mechanism. Systematic mutations and *in vivo* photo crosslinking experiments have previously suggested the functional role of the ExbD NIBS also in *E.coli*^29^, further emphasizing the importance of the opening and closing of ExbD as a general mechanism.

In the context of the full ExbB-ExbD complex, we hypothesize that the proton channel would predominantly occupy a main state with ExbD in a closed conformation, while a sparsely populated state would feature ExbD in an open conformation (Figure 2f). The closed state of ExbD would thus represent the inactive, impermeable state of the ExbB-ExbD proton channel, as it should be predominantly closed to prevent continuous proton leakage. Conversely, the open state of ExbD is thought to represent the active, proton-permeable state.

### The open state of ExbD is conformationally selected upon TonB binding

To explore the interaction between HasB/TonB and ExbD, we analyzed 2D [^1^H-^15^N]-TROSY spectra of ^15^N-labeled HasB^Sm, peri^ (residues 37-263) and ^15^N-labeled TonB^Ec, peri^ (residues 34-239) upon addition of their unlabeled partner ExbD^Sm, peri^ and ExbD^Ec, peri^ respectively (Figure 3a and Figure S11a).

**Figure 3.**
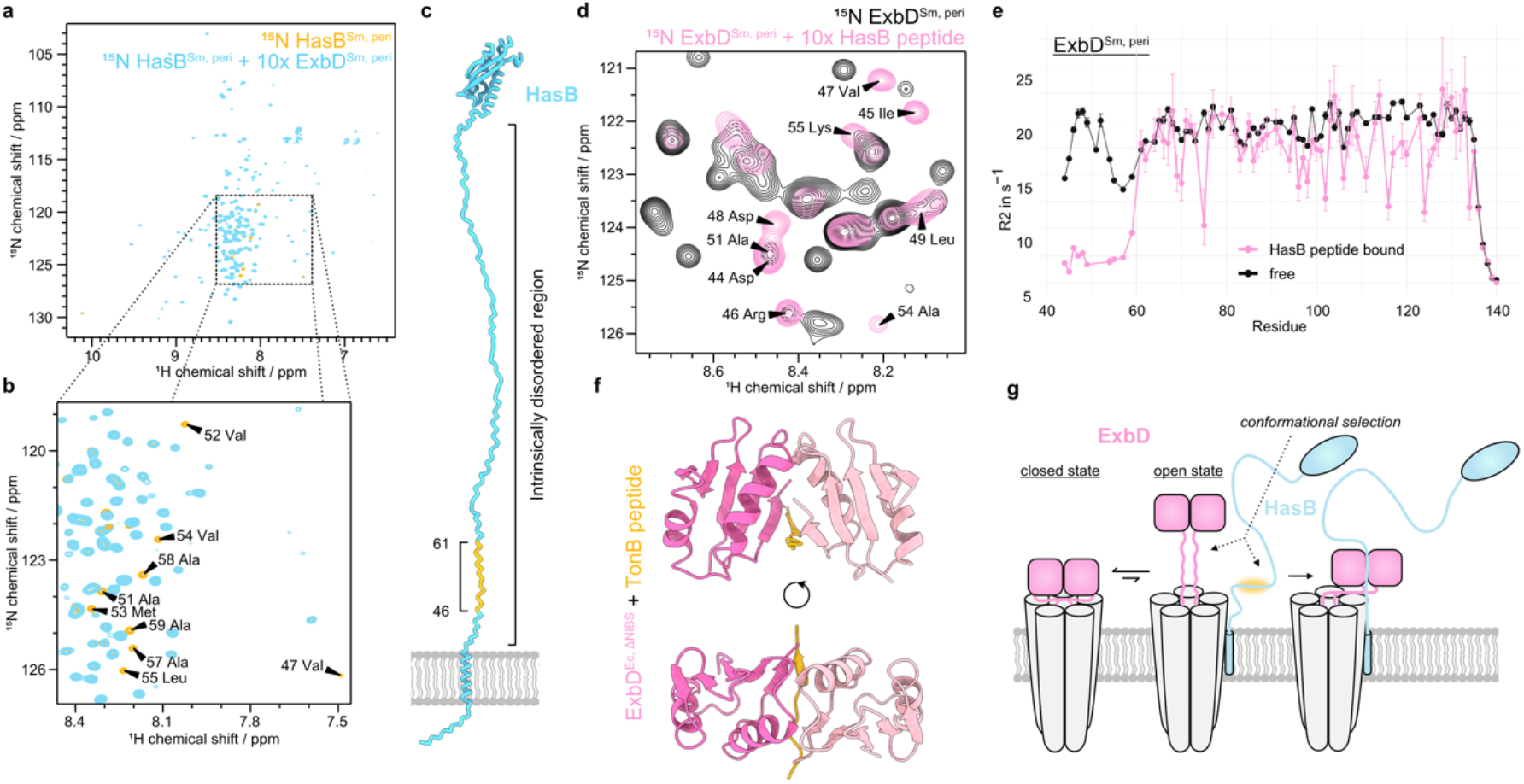
The open state of the periplasmic domain of ExbD is conformationally selected upon binding to TonB/HasB. **a,b** 2D [^1^H-^15^N]-TROSY spectra of ^15^N-labeled HasB^Sm, peri^ alone (orange) and in the presence of unlabeled ExbD^Sm, peri^ (blue). The presence of ExbD leads to the disappearance (broadening) of the peaks corresponding to the residues 46-61 of HasB^Sm, peri^ indicating binding of this region to ExbD. **c** Model of HasB inserted in the inner membrane: The HasB-ExbD region of interaction (orange) is located in an intrinsically disordered region (IDR) of HasB (blue) close to the inner membrane inserted N-terminal α-helix. **d** 2D [^1^H-^15^N]-HMQC spectra of ^15^N-labeled ExbD^Sm, peri^ without (black) and with (pink) the HasB peptide (corresponding to the binding region on the HasB side). Binding of the peptide leads to signal intensity decrease and appearance of new peaks that can be assigned to the NIBS residues. **f** R_2_ relaxation rate measurements of the free (black) and HasB peptide bound state (pink) of ^15^N ExbD^Sm, peri^ reveal that the binding stabilizes the open state of ExbD^Sm, peri^ characterized by the disordered nature of the NIBS residues (lower relaxation rates). This suggests a conformational selection of the open state of ExbD. **e** Deletion of the NIBS residues of ExbD^Ec, peri^ allows for crystallization with a peptide corresponding to the binding region on the TonB side. Herein, the TonB peptide (orange) replaces the NIBS residues at the interface in between the two subunits of the ExbD dimer (pink, light pink). **g** Schematic mechanism: The open state of ExbD (pink, right side) is conformationally selected by TonB/HasB (blue) leading to a disorder-to-order transition of the IDR of TonB/HasB. The binding motif on the TonB/HasB side is highlighted in orange.

A close examination of the central region of the spectra shows the disappearance of peaks corresponding to an N-terminal IDR of HasB and TonB (Figure 3b-c and Figure S11b-c), confirming their interaction with ExbD. To examine the changes on the ExbD side, we further analyzed 2D [^1^H-^15^N]-HMQC spectra of ^15^N-labeled ExbD^Sm, peri^ and ^15^N-labeled ExbD^Ec, peri^ with and without the respective HasB/TonB peptides that correspond to the binding regions. Peptide addition not only causes line broadening – indicative of interaction and conformational exchange between the peptide bound and free state – but also leads to the emergence of novel peaks in the center of the spectrum (Figure 3d and Figure S12). Interestingly, these new peaks can be assigned to the NIBS region, as they show the same chemical shifts as the NIBS residues in the minor, open conformation (State C) observed in the ^15^N-CEST experiment (Figure 2a). Additionally, low R_2_ relaxation rates confirm the disordered nature of the NIBS residues in the HasB-bound state (Figure 3e).

Herein, we initially hypothesized that TonB/HasB might bind between the two protomers of ExbD, effectively replacing the NIBS residues at the dimer interface. This idea was driven by our observation that the NIBS residues unfold upon addition of TonB/HasB, implying a potential competition between the NIBS residues and HasB/TonB for the same binding site. To investigate this possibility, we solved the crystal structure of an ExbD construct lacking the N-terminal residues 43-60 (ExbD^Ec, 1′NIBS^) in complex with the corresponding TonB peptide at 1.5 Å resolution. (Figure 3e and Table S5). The crystal structure confirms that the IDR of TonB undergoes a disorder-to-order transition upon binding to ExbD, replacing the NIBS residues in between the two ExbD protomers. In this arrangement, the TonB peptide forms an intermolecular β-sheet: A parallel β-sheet with the β6-strand of one protomer, and an antiparallel β-sheet with the β6-strand of the second one (Figure S13).

Our results support that in the context of the full ExbB-ExbD complex, ExbD exchanges between a closed state and a sparsely populated open state. Upon encountering HasB/TonB, this open state is conformationally selected through the binding of HasB/TonB into the groove between the two subunits, which was previously occupied by the now-unfolded NIBS (Figure 3g). This would mark a critical first step in the activation of the system.

### ExbD interacts transiently with the periplasmic peptidoglycan layer

Since HasB/TonB and ExbD reside in the periplasm in proximity to peptidoglycan, we explored the role of peptidoglycan in the system. For that purpose, we studied 2D [^1^H-^15^N] spectra of ^15^N-labeled ExbD^Sm, peri^ and ExbD^Ec, peri^ in the presence and absence of different peptidoglycan sacculi. Upon addition of *E. coli* peptidoglycan (Figure 4a and Figure S14) some peaks loose intensity, indicating the emergence of peptidoglycan-binding induced conformational exchange. Notably, this binding is transient since fully peptidoglycan-bound ExbD would not be visible by solution NMR spectroscopy due to the megadalton size of sacculi.

**Figure 4.**
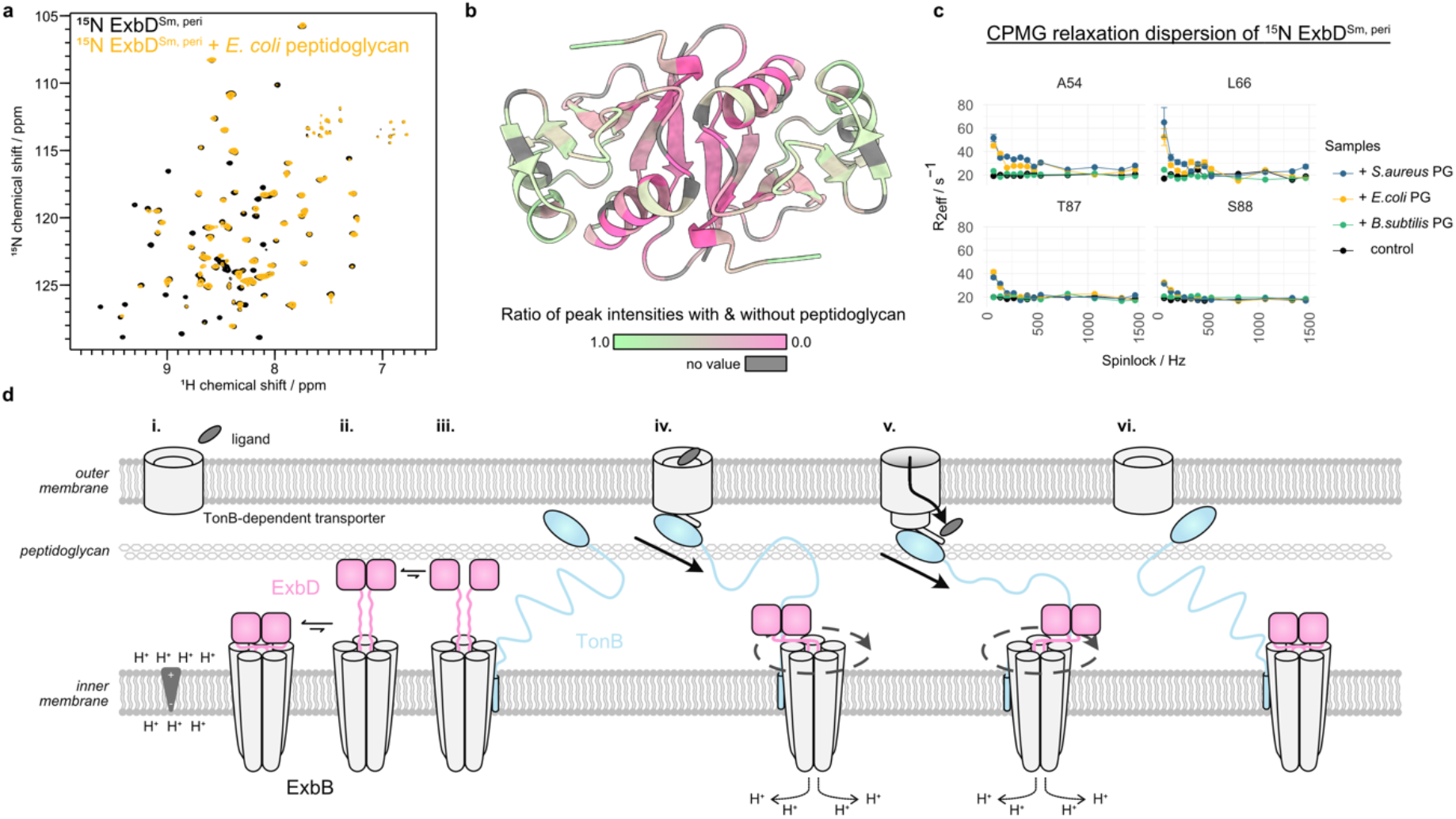
The periplasmic domain of ExbD interacts with the peptidoglycan layer. **a** 2D [^1^H-^15^N]-TROSY spectra of ^15^N ExbD^Sm, peri^ without (black) and with (orange) *E. coli* peptidoglycan. The interaction of ExbD^Sm, peri^ with peptidoglycan leads to the disappearance (broadening) of peaks. **b** Mapping this peak intensity loss in terms of a peak height ratio (pink to green, grey color means no value) onto the dimeric ExbD^Sm, peri^ structure shows that the interface within the dimer experiences the largest increase in exchange contribution to the R_2_ relaxation rate upon interaction with peptidoglycan. This suggests that possibly the dimer dissociates upon interaction. **c** This increase in R_2_ relaxation rate due to exchange can be measured by Car-Purcell-Meiboom-Gill (CPMG) relaxation dispersion experiments and reveals that ExbD^Sm, peri^ selectively interacts with peptidoglycan from *E.coli* (orange), *S.aureus* (blue) – they all show decaying profiles – but not *B. subtilis* (green), which shows the same flat curve as the control experiment without peptidoglycan (black). **d** Schematic representation of the full mechanism. In the resting state in the absence of ligand (i.), closed ExbD (pink) is in exchange with a minor open conformation (ii.). This extended, open conformation can reach the peptidoglycan layer, in interaction with which its dimer interface is loosened (iii.). This allows TonB/HasB (blue) to enter in between the ExbD monomers and to subsequently select the open state of ExbD by binding to it (iv.). Also, the C-terminal domain of TonB/HasB interacts with a ligand (grey) bound TonB-dependent transporter (TBDT). Proton motive force induced rotation of ExbD within ExbB leads to a pulling through the IDR of TonB/HasB opening the TBDT and enabling active transport of ligands (v.). Eventually, the pulling force becomes too large and leads to the dissociation of TonB/HasB from the TBDT, and the return to the resting state of the system (vi.).

Mapping the decrease of peak intensities of ExbD^Sm, peri^ onto the ExbD structure, shows that the dimeric interface is primarily affected by this exchange (Figure 4b). Consequently, we propose that upon binding to peptidoglycan the dimeric interface of ExbD is reorganized. This reorganization would require a prior dissociation of the dimer. Consistent with this, the disulfide-bridged ExbD^Sm, peri^ V47C mutant, which cannot undergo dimer dissociation, failed to interact with peptidoglycan (Figure S15). To pinpoint the interaction interface on the peptidoglycan side, we further studied the interaction of ExbD^Sm, peri^ with *Staphylococcus aureus* and *Bacillus subtilis* peptidoglycan. Compared to the peptidoglycan of *E. coli*, *S. aureus* possesses chemically different peptide stems whereas *B. subtilis* features glycans with a non-acetylated glucosamine moiety.^30^ Notably, among the peptidoglycans tested here, only the one from *B. subtilis* does not interact with ExbD^Sm, peri^, which pinpoints the interaction site on the peptidoglycan to the glycan chains (Figure S16). Additionally, peptidoglycan digested with mutanolysin, a muramidase, does not show any interaction with ExbD, further supporting this hypothesis (Figure S17). Furthermore, NMR relaxation dispersion experiments showed an increase in effective R_2_ relaxation rates upon the addition of peptidoglycans from *E. coli* and *S. aureus*, but not from *B. subtilis* (Figure 4c and Figure S18-19).

## DISCUSSION

We have elucidated the dimeric structure of the periplasmic domain of ExbD, which was previously unresolved in cryo-EM studies. The observed dimeric form is consistent with the two N-terminal, ExbB-integrated α1 helices of ExbD present in the cryo-EM maps. We have also demonstrated by NMR spectroscopy that the periplasmic domain of ExbD undergoes extensive conformational exchange on the millisecond timescale, sampling a primary closed state and at least one secondary open state with an unfolded NIBS region. This dynamic nature of the protein might explain its invisibility in the cryo-EM densities. The open state of ExbD is selectively bound by an IDR of the partner proteins TonB/HasB, which undergo a disorder-to-order transition as they bind between the two ExbD protomers, as evidenced by X-ray crystallography. Moreover, we have revealed that the periplasmic domain of ExbD selectively interacts with specific peptidoglycan species, most likely promoting dimer dissociation.

Taking our results together, we propose the following mechanism of action for the Ton system: A resting state is characterized by a unliganded TBDT (Figure 4d, stage i). Within the ExbB-ExbD complex, dimeric ExbD predominantly exists in a closed conformation and a sparsely populated open conformation (Figure 4d, stage ii.) as we have shown by NMR spectroscopy. The closed conformation likely represents the inactive, proton impermeable state of the proton channel. The sparsely populated open state of ExbD is triggered by the unfolding of the NIBS region, which subsequently reveals TonB/HasB-binding region between the ExbD protomers. Yet, even in this state, the TonB/HasB-binding region in between the ExbD dimer remains inaccessible. This is due to structural organization of the ExbD-ExbB complex: The N-terminal α1-helices of each protomer are inserted into the ExbB channel, and the C-terminal periplasmic domain of ExbD forms a tight dimer. However, the adaptation of the sparsely populated open state following the unfolding of the NIBS region brings ExbD into close proximity with the peptidoglycan layer. Here, ExbD binds to the glycan chains of the peptidoglycan. This binding event imposes the reorganization of the dimeric interface of ExbD and, hence, a dimer-to-monomer dissociation – permitting the penetration and interaction of TonB/HasB in between the ExbD protomers (Figure 4d, stage iii). The transient opening and the dynamics of the ExbD is thus a critical prerequisite for TonB/HasB binding and, consequently, the active transport of nutrients.

In the Tol-Pal system, dimeric TolR, a ExbD structurally homologous protein, was shown to form a complex with peptidoglycan. However, this complex formation is only possible when the β6-strand at the dimeric interface is removed yielding a different dimeric interface and organization, proposedly exposing a peptidoglycan binding motif.^24,31^ This is consistent with our observation that the peptidoglycan-interaction strongly affects the ExbD^Sm, peri^ dimer interface. However, in our case, the presence of β6 renders the interaction with peptidoglycan more transient as we see an exchange between bound and unbound ExbD (Figure 4a).

Upon ligand binding to the TBDT, its N-terminal extension containing the TonB box extends into the periplasm, recruiting the TonB protein. Intriguingly, molecular dynamics simulations of the IDR of TonB have shown that its Glu-Pro and Lys-Pro repeats can form a hairpin structure.^17^ This configuration may serve to sterically conceal the ExbD-binding motif of TonB. We hypothesize that with its anchoring points on ExbB within the inner membrane and on the TBDT in the outer membrane, the IDR of TonB undergoes tension as both anchoring points diffuse in their respective membranes. This tension is likely to destabilize the hairpin structure, thereby unmasking the binding motif. This newly accessible motif engages with the open state of ExbD, conformationally selecting its open state. This interaction effectively stabilizes the ExbB-ExbD complex in an active, proton-permeable state (Figure 4d, stage iv). However, in a single-target system such as the Has, the high affinity of HasB for the TBDT (nM K_d_ versus µM for TonB-TBDTs) suggests that HasB remains associated to the TBDT even in the absence of the ligand. Here, signaling of ligand-binding is proposedly transmitted via a stabilization of the HasB-TBDT interaction by involving more polar contacts, as shown in our previous study.^19^

After the stabilization of the ExbB-ExbD complex in the active state, the PMF is translated into a rotation of ExbD within ExbB through proton translocation via the de- and reprotonation of Asp 25 of ExbD (Figure 4d stage iv). This results in a wrapping of the intrinsically disordered linker of HasB/TonB around ExbD, exerting a pulling force on the TonB box of the TBDT. This force was shown to be sufficient in unfolding and removing the plug domain from the TBDT^32^, culminating in ligand import into the periplasm (Figure 4d, stage v). Finally, due to the exerted force, TonB/HasB dissociates from the TBDT. This marks the return to the resting state of the Ton system (Figure 4d, stage vi). This mechanism extends the wrap- and-pull model recently proposed by Ratliff *et al*.^18^

Our study uncovers a conserved interaction between HasB/TonB and the open state of ExbD in both single- and multi-target systems. However, we showed that the transition between the open and closed states of ExbD operates at distinct rates in these systems: on the slow NMR time scale (ms) for the single-target Has system and in the intermediate time scale (μs-ms) for the multi-target Ton system. This faster rate of conformational exchange in the *E. coli* Ton system may indicate a capacity for more rapid adaptation, an attribute beneficial for catering multiple TBDTs (up to 9 have been reported^33^), unlike the single-target HasB.

In homologues of ExbD of other PMF-dependent motors involved in membrane integrity (Tol-Pal system) and gliding motility (AglQRS system)^11^, the NIBS residues are highly conserved, suggesting a common mode of operation involving multiple states due to the unfolding of the NIBS. This extends to the ExbD-binding motif of TonB homologues, indicating a conserved mechanism of energy transduction from the proton channel to the outer membrane (Figure S20-21). We speculate that stabilizing the open state of ExbD or its homologues with a drug binding in between the protomers could not only inhibit the vital functioning of these motors but also lock the proton channel complexes into an active, proton permeable state, creating a fatal proton leakage in the inner membrane.

## METHODS

### Strains and plasmid construction

Strains, plasmids, and oligonucleotides are shown in Table S6. To produce ExbD mutants, the pBADexbBD_Sm_ or the pBAD24exbBDhis6_Sm_ plasmids were respectively amplified by PCR with oligonucleotides couples ExbDV475’/ ExbDV47C, ExbDV475’/ ExbDV47G, ExbDV475’/ ExbDV47R and ExbDE1115’/ ExbDD111C. The PCR product was digested with *Dpn*I, self-ligated and transformed in XL1-Blue. Recombinant clones were isolated and verified by sequencing.

### Protein expression and purification

Protein samples used in this study are summarized in Table S1. All ExbD variants and TonB^Ec, peri^ contain an N-terminal His-tag followed by a TEV cleavage site. The corresponding genes were synthesized and cloned by ProteoGenix into pet-30a(+) vectors, which were used to transform *E.coli* BL21 (DE3). For protein expression, glycerol stocks of transformed bacteria were used to inoculate 10 mL Luria-Bertani (LB) medium, which was grown for 8 h at 37 °C. In a second step, 100 mL M9 medium containing 4 g/L ^13^C-glucose and 1 g/L ^15^NH_4_Cl as the only carbon and nitrogen sources were added and the culture was incubated overnight at 30 °C. The next morning, 1 L fresh M9 medium was inoculated with the overnight culture to a starting optical density at 600 nm (OD) of 0.1. The bacterial cultures were grown to an OD of 0.7 at 37 °C and the expression was induced by the addition of 0.5 mM IPTG (isopropyl β-D-1-thiogalactopyranoside). Protein expression was conducted for 5 h at 37 °C (ExbD variants and TonB^Ec, peri^) or for 20 h at 30 °C (HasB_37-263, Sm_). For non-NMR studies, all protein expression steps were conducted in LB medium instead of M9 medium. The bacterial cells were harvested by centrifugation at 7000xg for 15 min at 37 °C, resuspended in Buffer A (20 mM sodium phosphate, 30 mM imidazole, 500 mM NaCl, pH 7.4) containing one EDTA-free protease inhibitor pill (Roche) per 25 mL and stored at −80 °C.

For expression of deuterated protein used for the detection of hydrogen bonds by NMR spectroscopy, bacteria were adapted to and grown in fully deuterated M9 medium containing ^13^C,D_7_-glucose and ^15^ND_4_Cl as the only carbon and nitrogen sources.

For protein purification, the bacterial cell pellets were thawed and 1.5 μL benzonase (Millipore) were added per 25 mL of cell suspension to digest nucleic acids. Cells were disrupted by sonication with a Vibracell 72405 sonicator (2s on, 1 s off, 80% amplitude and a total time of 20 min). Insoluble cell debris was removed from the soluble protein-containing supernatant by centrifugation at 30.000xg at 4 °C for 30 min and by sterile filtration through a 0.22 μm membrane. By means of an Akta system, the lysate was loaded onto a pre-equilibrated (Buffer A: 20 mM sodium phosphate, 30 mM imidazole, 500 mM NaCl, pH 7.4) nickel affinity chromatography column (two stacked 5 mL HisTrap Chelating HP, Cytiva), washed with 5 column volumes (CV) Buffer A and eluted with a linear gradient of 0-100 % Buffer B (20 mM sodium phosphate, 500 mM imidazole, 500 mM NaCl, pH 7.4) over 5 CV. The protein containing fractions as judged by SDS-PAGE were united and diluted (at least 10x) in TEV buffer (50 mM Tris-HCl, 1 mM dithiothreitol, pH 8.0). 1 mg of TEV protease was added per 100 mg of target protein and cleavage of the His-tag was conducted at 34 °C for 2 h. Then the mixture was loaded onto a HisTrap HP column to remove TEV-His_6_ and un-cleaved proteins. The target protein containing fractions from the washing step were pooled, concentrated to about 1.5 mL and injected with a 2 mL loop onto a pre-equilibrated (Buffer C: 50 mM sodium phosphate, 50 mM NaCl, pH 7.0) size-exclusion chromatography column (HiLoad 16/600 Superdex 75 pg, Cytiva or HiPrep 16/60 Sephacryl S-100 HR, Cytiva) and eluted with 1.5 CV Buffer C. The fractions containing the target protein were pooled, aliquoted, flash-frozen in liquid nitrogen and stored at −20 °C. For X-ray crystallography sample preparation, Buffer D (50 mM Tris-HCl, 50 mM NaCl, pH 7.0) was used for size-exclusion chromatography.

For the His-tag lacking protein HasB^Sm, peri^, the nickel affinity column steps were replaced by a cation-exchange chromatography step. For that purpose, bacterial cells were resuspended in Buffer E (50 mM Tris HCl, 100 mM NaCl, pH 8.8) after harvest. For purification, the disrupted lysate was loaded onto a pre-equilibrated (Buffer E: 50 mM Tris HCl, 100 mM NaCl, pH 8.8) cation-exchange chromatography column (SP Sepharose HP 16/60, Cytiva), washed with 10 CV Buffer E and eluted with a linear gradient of 0-100 % Buffer F (50 mM Tris HCl, 1000 mM NaCl, pH 8.8) over 15 CV. The protein containing fractions as judged by SDS-PAGE were united, concentrated and submitted to size-exclusion chromatography as mentioned above.

To acquire intermolecular NMR restraints, a sample containing 50% ^15^N,^13^C-ExbD^Sm, peri^ and 50% ^14^N,^12^C-ExbD^Sm, peri^ (50% labeled – 50% unlabeled) was prepared. For that purpose, the two batches of purified protein were mixed in equimolar quantities, denatured in 8 M Urea and subsequently refolded by dialyzing to Buffer C (3x 1 L Buffer C). Fingerprint NMR spectra showed no difference between native and refolded proteins.

### Analytical ultracentrifugation (AUC)

For sedimentation velocity experiments, ExbD^Sm, peri^ and ExbD^Ec, peri^ at concentrations of 20 μM, 100 μM and 1000 μM were loaded into 3 mm or 1.2 cm centerpieces and centrifuged overnight at 42000 rpm in a Beckman Coulter Optima AUC centrifuge operating with an AN60-Ti rotor. Fitting of the data using a continuous size distribution c(S) model was conducted via SEDFIT 15.1 with a confidence level (F-ratio) of 0.95. From the fit, sedimentation coefficients at zero concentration in the buffer (50 mM sodium phosphate, 50 mM NaCl, pH 7.0) could be calculated as well as molecular weights estimated. ^34^ Hydrodynamic parameters of the closed-state ExbD^Sm, peri^ were calculated with Hydropro.^35^ For the closed-state the NMR-structure determined in this study was used, the sedimentation coefficient of monomeric ExbD was calculated from PDB ID 2PFU.^10^

### Solution NMR spectroscopy and resonance assignment

All used NMR samples are summarized in Table S2. All spectra were acquired using Topspin4 on either a 600 MHz Avance III HD or a 800 MHz Avance NEO spectrometer equipped with cryogenically cooled triple resonance ^1^H[^13^C/^15^N] probes (Bruker Biospin). For pulse calibration and setting up standard experiments NMRlib was used.^36^ For spectral fingerprinting, either 2D ^15^N-^1^H SOFAST-HMQC or BEST-TROSY spectra were recorded.^37^ For backbone assignments, generally 3D BEST-HNCA/-HNcoCA/-HNCO/-HNCACB/-HNcoCACB and BEST-TROSY HNcaCO spectra were acquired with 25% non-uniform sampling (NUS).^38^ Sidechain assignments were conducted by the acquisition of 3D HBHAcoNH and HCCH-TOCSY as well as 2D hbCBcgcdHD and 2D hbCBcgcdceHE spectra. For structural restraints, 3D ^15^N-edited NOESY-HSQC, ^13^C^aromatic^-edited NOESY-HSQC and ^13^C^aliphatic^-edited NOESY-HSQC, and, for intermolecular structural restraints, ^13^C-,^15^N-filtered 3D NOESY-^13^C-HSQC and ^13^C-,^15^N-filtered 3D NOESY-^15^N-HSQC spectra were recorded at a B_0_ field strength of 800 MHz. A mixing time of 120 ms was used for all NOESY spectra. For the detection of hydrogen bonds, a 3D BEST-TROSY HNCO spectrum with a total N-CO INEPT transfer period of 133 ms was acquired.^39^ Spectral data were referenced to 2,2-Dimethyl-2-silapentane-5-sulfonate (DSS), processed with NMRPipe^40^ and in the case of NUS reconstructed using SMILE^41^. Visualization of spectra, peak picking and resonance assignments were conducted using CcpNmr version 2.5^42^ and version 3^43^.

The final resonance assignment of ExbD^Sm, peri^ was deposited to the Biological Magnetic Resonance Data Bank (BMRB) under the accession code 34826.

^15^N-R_1_ and -R_2_ relaxation rates were measured by pseudo-3D experiments (^1^H, delay, ^15^N). For ^15^N-R_1_ measurements the relaxation delays were 0, 20 70, 150, 200, 300, 400, 550, 700, 900, 1100, 1500, 1800 and 2000 ms in random order with a recycle delay of 5.5 s between scans. For ^15^N-R_2_ measurements the relaxation delays were 0, 66.67, 133.33, 200, 266.67, 400, 533.33, 800, 1066.67, 1333.33 and 1466.67 ms in random order with a recycle delay of 4.5 s between scans. Peak heights and errors were extracted with the non-linear N-dimensional spectral modeling (nlinLS) program as part of the NMRPipe suite^40^, and fitted to a monoexponential function. Hereby, the errors were estimated by running 21 Monte Carlo trials for each fit. For ^15^N[^1^H] heteronuclear nuclear Overhauser effect (nOe) measurements, two HSQC spectra without (reference) and with a ^1^H saturation pulse (1 ms, 250 Hz), and with a pre-saturation delay of 3 s and a recycle delay of 10.5 s were recorded. Peak heights and errors were extracted from the two spectra with nlinLS and heteronuclear nOe values calculated as the ratio of peak heights with and without saturation pulse.

Carr-Purcell-Meiboom-Gill (CPMG) relaxation dispersion profiles were measured by BEST-TROSY pseudo-3D experiments (^1^H, ^15^N spinlock strength, ^15^N) at a B_0_ field strength of 600 and 800 MHz. For these experiments the spinlock strengths were 0, 66.67, 133.33, 200, 266.67, 400, 800, 1066.67, 1333.33 and 1466.67 Hz in random order during a total relaxation delay T_relax_ of 30 ms. Peak heights and errors were extracted from the spectral data with nlinLS and used to calculate the spinlock dependent R_2eff_ rate following equation 1.

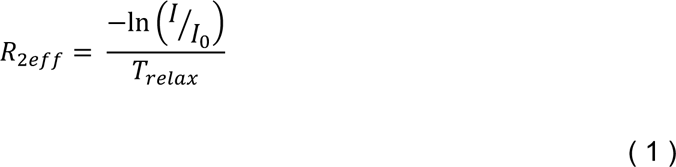

The spinlock dependent R_2eff_ rates were fitted numerically to a two-state exchange with ChemEx^28^ by minimizing the target function *χ*^2^ (See equation 2).

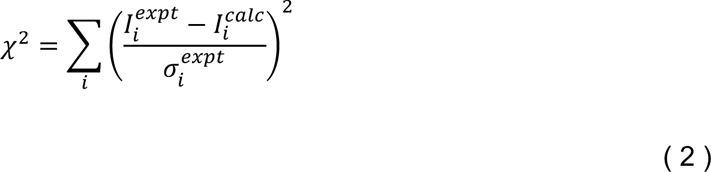

Chemical exchange saturation transfer (CEST) profiles were measured by two pseudo-3D experiments (^1^H, ^15^N carrier position, ^15^N) with ^15^N spinlock strengths of 22.7 Hz and 55.9 Hz applied for a duration T_ex_ of 400 ms at a B_0_ field strength of 800 MHz. The CEST dimension had in both cases a spectral width of 2854.25 Hz, and for 22.7 Hz spinlock strength increments of 20.3 Hz and for 55.9 Hz spinlock strength increments of 40.6 Hz. The exact spinlock strength was determined by an incremented nutation experiment.^44^ Peak heights and errors were extracted from the spectral data with nlinLS, and fitted to a three-state exchange with ChemEx by minimizing the target function *χ*^2^ (See equation 2). The errors were calculated based on the scatter observed in the CEST profiles using the ChemEx function “scatter”.

For the Deuterium-Hydrogen exchange experiments, we used a ExbD^Sm, peri^ sample that was previously exchanged to D_2_O After 2 weeks, this sample was lyophilized overnight, then redissolved in Buffer C (50 mM sodium phosphate, 50 mM NaCl, pH 7.0). The first SO-FAST HMQC was recorded after 2 min 50 s with subsequent 99 time points of 2 min and 39 s. A final reference spectrum was recorded after 24 h. The peak intensities were extracted with nlinLS and fitted with fitXY.tcl of the NMRpipe suite^40^ to the complement of an exponential decay of the form F(t)=A[1-e^(k*t)], where A is a normalization factor depending on the maximal peak height, t the exchange time and k the exchange rate.

For examining the interaction between ExbD protein species and peptidoglycan, peptidoglycan sacculi from *E. coli*, *B. subtilis* or *S. aureus* (InvivoGen) were washed twice with Buffer C and added to a 100 μM protein-containing NMR sample. Simultaneously, a reference sample – maintaining the same protein concentration but devoid of peptidoglycan – was also prepared.

### NMR structure calculation of the periplasmic domain of ExbD

NMR structure calculation was performed by the ARIAweb server using the standard torsion angle simulating annealing protocol.^45^ Firstly, the unambiguous intermolecular NMR restraints were used to assign peaks in the 3D ^15^N-edited NOESY-HSQC, ^13^C^aromatic^-edited NOESY-HSQC and ^13^C^aliphatic^-edited NOESY-HSQC spectra. The rationale behind this was that peak lists of the intermolecular NMR restraints cannot directly be used during the structure calculation protocol because their sparsity makes them un-calibratable. Additionally, further unambiguous peaks in the NOESY spectra were assigned. Torsion angles were predicted based on assigned chemical shift values using TALOS-N.^46^ Secondary structure information was also used as “structural rules” in ARIA to initially prevent incorrect assignments of intermolecular NOE as intramolecular. The CcpNmr project file containing all assigned chemical shifts, NOESY peak lists and hydrogen bonds was uploaded to the ARIAweb server. Here, 9 iterations of automatic peak assignment and calibration of the 3D ^15^N-edited NOESY-HSQC, ^13^C^aromatic^-edited NOESY-HSQC and ^13^C^aliphatic^-edited NOESY-HSQC spectra was conducted in the following way: Manual assignments were used and flagged as reliable, diagonal peaks were filtered, and structural rules enabled. The error of chemical shifts based on the resonance assignment was used in addition to a ^1^H chemical shift tolerance of 0.04/0.02 ppm in the indirect/direct dimensions and a ^13^C/^15^N chemical shift tolerance of 0.3 ppm. For spin diffusion correction, a molecule correlation time of 10 ns was adopted. During structure calculation, a C2 symmetry was applied along with NCS restraints^47^, where the latter were used only until iteration 4. Dihedral angles derived from the TALOS-N predictions were supplied together with hydrogen bonds. In all 9 iterations 50 structures were calculated and the 7 best structures (based on their total energy) were used for analysis. Torsion angle simulating annealing was performed with 30000 high-temperature steps, 30000 cool1 steps and 30000 cool2 steps, and a log-harmonic potential was applied using automatic restraints weighting during the cool2 step of the simulated annealing.^48^ The 10 best structures of the last iteration were refined in an explicit shell of water molecules.^49^ The structure calculation protocol was iteratively repeated and some manual changes were applied to the NOE assignments in between runs. In the end, a consensus structure was calculated following the methodology reported.^50^ Briefly, 20 independent ARIA runs were carried out using the same input data but different random number seeds, generating varying random initial conformations and velocities for the molecular dynamics simulated annealing protocol. Subsequently, cross-peaks that remained active (i.e., those retaining at least one assignment possibility) were collected at the conclusion of 12 out of the 20 ARIA runs. For each active cross-peak, assignment possibilities from each individual ARIA run were combined to generate a new list of consensus distance restraints. Lastly, a single iteration ARIA run was conducted using the consensus distance restraints as input, resulting in the final consensus structure ensemble.

The final structural ensemble of ExbD^Sm, peri^ and the restraints were deposited to the Protein Data Bank (PDB) under the accession code 8PEK.

### Bacterial growth test

Strain *E. coli* K12 C600*ΔhemAΔexbBDΔtonB*(pAMhasISRADEB) was transformed with plasmid pBADexbBD_Sm_ or its derivatives bearing mutation in *exbD*. A few colonies were first inoculated in 3 ml of LB medium at 37°C with the corresponding antibiotics, and 40 µM dipyridyl, 4µg/ml arabinose. Once the culture reached an OD_600nm_ of ca. 1.2-1.5, it was diluted and inoculated in 48 well Greiner plates, in the same medium to which was added 1 µM He-BSA, as a heme source. The initial OD_600nm_ of the cultures was 0.001. Each well contained 300 µl of growth medium. Triplicates of each strain were made, and the plate was incubated at 37°C with vigorous shaking (500 rpm) in a Clariostar Plus Microplate reader. OD_600nm_ was recorded every 15 minutes for 60 hours.

### Isolation of the ExbBD _Sm_ complex

Strain *E. coli* K12 C600*ΔhemAΔexbBDΔtonB*(pAMhasISRADEB) was transformed with the pBAD24exbBDhis6_Sm_ plasmid or its variants pBAD24exbBDV47Chis6_Sm_, pBAD24exbBDV47Ghis6_Sm_, pBAD24exbBDV47Rhis6_Sm_, and pBAD24exbBDD111Chis6_Sm_. For the expression of each variant, 200 ml of culture in LB at 37°C (in the presence of ampicillin (100 µg/ml), spectinomycin (75 µg/ml), delta-aminolevulinic acid (25µg/ml) was induced with 40 µg/ml arabinose at OD_600nm_ of 0.5 and the culture continued for 3 hours. Cells were harvested, and the pellet washed and resuspended in 100 mM Tris-HCl pH 8.0, 1 mM EDTA, in the presence of lysozyme (50 µg/ml); after one freeze-thaw cycle, the suspension was sonicated, MgSO_4_ was added at a final concentration of 4 mM, DNase was added, and the suspension centrifuged at 16000 g for 45 minutes. The pellet was resuspended, and solubilized in 20 mM Tris-HCl pH 8.0, 100 mM NaCl, 20 mM Imidazole, 10% glycerol, 0,8% LMNG, at a final concentration of 160 OD_600nm_ equivalent/ml. After 45 minutes, the suspension was centrifuged (16000 g, 45 minutes), and the supernatant incubated with Ni-Agarose. After 3 hrs, of incubation, the beads were washed 4 times with 20 mM Tris-HCl pH 8.0, 100 mM NaCl, 20 mM Imidazole, 10% glycerol, 0,005% LMNG, the bound proteins eluted in the same buffer in the presence of 200 mM Imidazole. The equivalent of 8.5 OD_600nm_ was loaded on each gel lane.

### Crystallization and diffraction collection

For the structural analysis of the ExbD–TonB complex, 250 μL of 2 mM (20 mg/mL) ExbD ^Ec, 1′NIBS^ with 10 mM TonB peptide (KKAQPISVTMVTPADLEPPQAKK) in Buffer G (50 mM Tris-HCl, 50 mM NaCl, pH 7.0) were prepared. Firstly, preliminary screening of crystallization conditions was performed using the vapor diffusion technique with a MosquitoTM nanoliter-dispensing system (TTP Labtech, Melbourn, United Kingdom) in accordance with established protocols.^51^ Specifically, sitting drops were prepared by combining 400 nL of protein-peptide mixture with crystallization solutions (comprising 672 commercially available conditions) in a 1:1 ratio and then equilibrating the mixture against a 150-μL reservoir in 96-well plates (Greiner Bio-one, GmbH, Frichenhausen, Germany). The crystallization plates were subsequently maintained at 18 °C in an automated RockImager1000 (Formulatrix, Bedford, MA, United States) imager to monitor the growth of crystals. The best crystals were obtained in 10% w/v glycerol and 3 M ammonium sulfate. The crystals were cryoprotected by soaking them into the crystallization solution supplemented with 20% glycerol as a cryoprotectant before freezing in liquid nitrogen. Subsequently, diffraction data were acquired at cryogenic temperatures (100 K) on the PROXIMA-1 beamline at the SOLEIL synchrotron facility (St Aubin, France) and processed with XDS^52^ through XDSME (https://github.com/legrandp/xdsme)^53^.

### X-ray structure determination and refinement

AlphaFold2 multimer^54^ was used to generate initial models of the ExbD-TonB peptide complex. On that basis, crystal structures were determined by molecular replacement with Phaser^55^. The final models were generated via an iterative process that involved manual model building using Coot^56^ and refinement in reciprocal space with REFMAC^57^ and Phenix^58^ All data collection details as well as model refinement statistics are summarized in Table S3.

The ExbD-TonB peptide complex structure was deposited to the Protein Data Bank (PDB) under the accession code 8P9R.

### Digestion of peptidoglycan

Five mg *E. coli* peptidoglycan (InvivoGen) were washed twice with Buffer C and finally resuspended in 800 μL. An aliquot of 25 μL of a 5 mg/mL stock solution of mutanolysin (Sigma Aldrich) were added and the mixture was incubated for 2 days at 37 °C. Considering fully glycosidically-digsted non-crosslinked peptidoglycan (GlcNAc-MurNAc-L-Ala-D-Glu-DAP-D-Ala; molecular weight = 2060 g/mol), this yields a final concentration estimated at 2.94 mM peptidoglycan fragments. The enzymatic reaction was terminated by heating the mixture at 95 °C for 10 min. After this, the mixture was filtered through a filter with a 10 kDa cutoff to remove larger peptidoglycan fragments and any remaining mutanolysin.

### Figure creation

All depictions of protein structures were generated with ChimeraX.^59^

## ACKNOWLEDGEMENTS

This work was supported by the French Agence Nationale de la Recherche (ANR Energir ANR-21-CE11-0039), the INCEPTION program « Investissement d’Avenir grant ANR-16-CONV-0005 and the Equipex CACSICE (ANR-11-EQPX-0008). We acknowledge NMR, Biophysical and Crystallography platforms of C2RT at the Institut Pasteur for their help and assistance. The 800-MHz NMR spectrometer and the optima AUC of the Institut Pasteur were partially funded by the Région Ile de France (SESAME 2014 NMRCHR grant no 4014526) and DIM one health, respectively. We thank Michael Nilges for his constant support and fruitful discussions. Molecular graphics and analyses were performed with UCSF ChimeraX, developed by the Resource for Biocomputing, Visualization, and Informatics at the University of California, San Francisco, with support from National Institutes of Health R01-GM129325 and the Office of Cyber Infrastructure and Computational Biology, National Institute of Allergy and Infectious Diseases.

## AUTHOR CONTRIBUTIONS

N.I-P and M.Z conceived the study. M.Z. produced the protein samples and performed NMR measurements. M.L. assisted in the wet lab. M.Z. and N.I-P conceived and analyzed NMR experiments. M.Z. and B.B. performed the NMR structure calculation. A.M. solved the X-ray structure. I.G.B helped in peptidoglycan data analysis. P.D. conducted the *in vivo* experiments. M.Z. designed the figures and wrote the manuscript with contributions of all authors. N.I-P. guided in the writing process. All authors approved the manuscript.

## REFERENCES

1. Blair, J. M. A., Webber, M. A., Baylay, A. J., Ogbolu, D. O. & Piddock, L. J. V. Molecular mechanisms of antibiotic resistance. Nat. Rev. Microbiol. 13, 42–51 (2015).

2. Schauer, K., Rodionov, D. A. & de Reuse, H. New substrates for TonB-dependent transport: do we only see the ‘tip of the iceberg’? Trends Biochem. Sci. 33, 330–338 (2008).

3. Noinaj, N., Guillier, M., Barnard, T. J. & Buchanan, S. K. TonB-Dependent Transporters: Regulation, Structure, and Function. Annu. Rev. Microbiol. 64, 43–60 (2010).

4. Paquelin, A., Ghigo, J. M., Bertin, S. & Wandersman, C. Characterization of HasB, a Serratia marcescens TonB-like protein specifically involved in the haemophore-dependent haem acquisition system. Mol. Microbiol. 42, 995–1005 (2001).

5. Biou, V. et al. Structural and molecular determinants for the interaction of ExbB from Serratia marcescens and HasB, a TonB paralog. *Commun*. Biol. 5, 1–15 (2022).

6. Amorim, G. C. de et al. The Structure of HasB Reveals a New Class of TonB Protein Fold. PLOS ONE 8, e58964 (2013).

7. Celia, H. et al. Cryo-EM structure of the bacterial Ton motor subcomplex ExbB–ExbD provides information on structure and stoichiometry. *Commun*. Biol. 2, 1–6 (2019).

8. Celia, H. et al. Structural insight into the role of the Ton complex in energy transduction. Nature 538, 60–65 (2016).

9. Maki-Yonekura, S. et al. Hexameric and pentameric complexes of the ExbBD energizer in the Ton system. eLife 7, e35419 (2018).

10. Garcia-Herrero, A., Peacock, R. S., Howard, S. P. & Vogel, H. J. The solution structure of the periplasmic domain of the TonB system ExbD protein reveals an unexpected structural homology with siderophore-binding proteins. Mol. Microbiol. 66, 872–889 (2007).

11. Rieu, M., Krutyholowa, R., Taylor, N. M. I. & Berry, R. M. A new class of biological ion-driven rotary molecular motors with 5:2 symmetry. Front. Microbiol. 13, (2022).

12. Deme, J. C. et al. Structures of the stator complex that drives rotation of the bacterial flagellum. Nat. Microbiol. 5, 1553–1564 (2020).

13. Shultis, D. D., Purdy, M. D., Banchs, C. N. & Wiener, M. C. Outer Membrane Active Transport: Structure of the BtuB:TonB Complex. Science 312, 1396–1399 (2006).

14. Josts, I., Veith, K. & Tidow, H. Ternary structure of the outer membrane transporter FoxA with resolved signalling domain provides insights into TonB-mediated siderophore uptake. eLife 8, e48528 (2019).

15. Domingo Köhler, S., Weber, A., Howard, S. P., Welte, W. & Drescher, M. The proline-rich domain of TonB possesses an extended polyproline II-like conformation of sufficient length to span the periplasm of Gram-negative bacteria. Protein Sci. 19, 625–630 (2010).

16. Evans, J. s., Levine, B. a., Trayer, I. p., Dorman, C. j. & Higgins, C. f. Sequence-imposed structural constraints in the TonB protein of E. coli. FEBS Lett. 208, 211–216 (1986).

17. Virtanen, S. I., Kiirikki, A. M., Mikula, K. M., Iwaï, H. & Ollila, O. H. S. Heterogeneous dynamics in partially disordered proteins. Phys. Chem. Chem. Phys. 22, 21185–21196 (2020).

18. Ratliff, A. C., Buchanan, S. K. & Celia, H. The Ton Motor. Front. Microbiol. 13, (2022).

19. Lefèvre, J., Delepelaire, P., Delepierre, M. & Izadi-Pruneyre, N. Modulation by Substrates of the Interaction between the HasR Outer Membrane Receptor and Its Specific TonB-like Protein, HasB. J. Mol. Biol. 378, 840–851 (2008).

20. Kaserer, W. A. et al. Insight from TonB Hybrid Proteins into the Mechanism of Iron Transport through the Outer Membrane. J. Bacteriol. 190, 4001–4016 (2008).

21. Kojima, S. et al. The Helix Rearrangement in the Periplasmic Domain of the Flagellar Stator B Subunit Activates Peptidoglycan Binding and Ion Influx. Structure 26, 590–598.e5 (2018).

22. Szczepaniak, J. et al. The lipoprotein Pal stabilises the bacterial outer membrane during constriction by a mobilisation-and-capture mechanism. Nat. Commun. 11, 1305 (2020).

23. Parsons, L. M., Lin, F. & Orban, J. Peptidoglycan Recognition by Pal, an Outer Membrane Lipoprotein,. Biochemistry 45, 2122–2128 (2006).

24. Wojdyla, J. A. et al. Structure and Function of the Escherichia coli Tol-Pal Stator Protein TolR*. J. Biol. Chem. 290, 26675–26687 (2015).

25. Hantke, K. & Braun, V. Functional Interaction of the tonA/tonB Receptor System in Escherichia coli. J. Bacteriol. 135, 190–197 (1978).

26. Jumper, J. et al. Highly accurate protein structure prediction with AlphaFold. Nature 596, 583–589 (2021).

27. Vallurupalli, P., Bouvignies, G. & Kay, L. E. Studying “Invisible” Excited Protein States in Slow Exchange with a Major State Conformation. J. Am. Chem. Soc. 134, 8148–8161 (2012).

28. Bouvignies, G. ChemEx.

29. Kopp, D. R. & Postle, K. The Intrinsically Disordered Region of ExbD Is Required for Signal Transduction. J. Bacteriol. 202, e00687–19 (2020).

30. Moynihan, P. J., Sychantha, D. & Clarke, A. J. Chemical biology of peptidoglycan acetylation and deacetylation. Bioorganic Chem. 54, 44–50 (2014).

31. Parsons, L. M., Grishaev, A. & Bax, A. The Periplasmic Domain of TolR from Haemophilus influenzae Forms a Dimer with a Large Hydrophobic Groove: NMR Solution Structure and Comparison to SAXS Data,. Biochemistry 47, 3131–3142 (2008).

32. Gumbart, J., Wiener, M. C. & Tajkhorshid, E. Mechanics of Force Propagation in TonB-Dependent Outer Membrane Transport. Biophys. J. 93, 496–504 (2007).

33. Grinter, R. & Lithgow, T. The structure of the bacterial iron–catecholate transporter Fiu suggests that it imports substrates via a two-step mechanism. J. Biol. Chem. 294, 19523–19534 (2019).

34. Schuck, P. Size-Distribution Analysis of Macromolecules by Sedimentation Velocity Ultracentrifugation and Lamm Equation Modeling. Biophys. J. 78, 1606–1619 (2000).

35. Ortega, A., Amorós, D. & García de la Torre, J. Prediction of Hydrodynamic and Other Solution Properties of Rigid Proteins from Atomic- and Residue-Level Models. Biophys. J. 101, 892–898 (2011).

36. Favier, A. & Brutscher, B. NMRlib: user-friendly pulse sequence tools for Bruker NMR spectrometers. J. Biomol. NMR 73, 199–211 (2019).

37. Schanda, P., Kupče, Ē. & Brutscher, B. SOFAST-HMQC Experiments for Recording Two-dimensional Deteronuclear Correlation Spectra of Proteins within a Few Seconds. J. Biomol. NMR 33, 199–211 (2005).

38. Lescop, E., Schanda, P. & Brutscher, B. A set of BEST triple-resonance experiments for time-optimized protein resonance assignment. J. Magn. Reson. 187, 163–169 (2007).

39. Cordier, F., Nisius, L., Dingley, A. J. & Grzesiek, S. Direct detection of N−H⋯O=C hydrogen bonds in biomolecules by NMR spectroscopy. Nat. Protoc. 3, 235–241 (2008).

40. Delaglio, F. et al. NMRPipe: A multidimensional spectral processing system based on UNIX pipes. J. Biomol. NMR 6, 277–293 (1995).

41. Ying, J., Delaglio, F., Torchia, D. A. & Bax, A. Sparse multidimensional iterative lineshape-enhanced (SMILE) reconstruction of both non-uniformly sampled and conventional NMR data. J. Biomol. NMR 68, 101–118 (2017).

42. Vranken, W. F. et al. The CCPN data model for NMR spectroscopy: Development of a software pipeline. Proteins Struct. Funct. Bioinforma. 59, 687–696 (2005).

43. Skinner, S. P. et al. CcpNmr AnalysisAssign: a flexible platform for integrated NMR analysis. J. Biomol. NMR 66, 111–124 (2016).

44. Guenneugues, M., Berthault, P. & Desvaux, H. A Method for DeterminingB1Field Inhomogeneity. Are the Biases Assumed in Heteronuclear Relaxation Experiments Usually Underestimated? J. Magn. Reson. 136, 118–126 (1999).

45. Allain, F., Mareuil, F., Ménager, H., Nilges, M. & Bardiaux, B. ARIAweb: a server for automated NMR structure calculation. Nucleic Acids Res. 48, W41–W47 (2020).

46. Shen, Y. & Bax, A. Protein backbone and sidechain torsion angles predicted from NMR chemical shifts using artificial neural networks. J. Biomol. NMR 56, 227–241 (2013).

47. Bardiaux, B. et al. Influence of different assignment conditions on the determination of symmetric homodimeric structures with ARIA. Proteins Struct. Funct. Bioinforma. 75, 569–585 (2009).

48. Nilges, M. et al. Accurate NMR Structures Through Minimization of an Extended Hybrid Energy. Structure 16, 1305–1312 (2008).

49. Linge, J. P., Williams, M. A., Spronk, C. A. E. M., Bonvin, A. M. J. J. & Nilges, M. Refinement of protein structures in explicit solvent. Proteins Struct. Funct. Bioinforma. 50, 496–506 (2003).

50. Buchner, L. & Güntert, P. Systematic evaluation of combined automated NOE assignment and structure calculation with CYANA. J. Biomol. NMR 62, 81–95 (2015).

51. Weber, P. et al. High-Throughput Crystallization Pipeline at the Crystallography Core Facility of the Institut Pasteur. Molecules 24, 4451 (2019).

52. Kabsch, W. XDS. Acta Crystallogr. D Biol. Crystallogr. 66, 125–132 (2010).

53. Legrand, P. XDSME: XDS Made Easier. (2017).

54. Evans, R. et al. Protein complex prediction with AlphaFold-Multimer. 2021.10.04.463034 Preprint at https://doi.org/10.1101/2021.10.04.463034 (2022).

55. McCoy, A. J. et al. Phaser crystallographic software. J. Appl. Crystallogr. 40, 658–674 (2007).

56. Emsley, P. & Cowtan, K. Coot: model-building tools for molecular graphics. Acta Crystallogr. D Biol. Crystallogr. 60, 2126–2132 (2004).

57. Murshudov, G. N. et al. REFMAC5 for the refinement of macromolecular crystal structures. Acta Crystallogr. D Biol. Crystallogr. 67, 355–367 (2011).

58. Liebschner, D. et al. Macromolecular structure determination using X-rays, neutrons and electrons: recent developments in Phenix. Acta Crystallogr. Sect. Struct. Biol. 75, 861–877 (2019).

59. Goddard, T. D. et al. UCSF ChimeraX: Meeting modern challenges in visualization and analysis. Protein Sci. 27, 14–25 (2018).

